# Charge transfer of unconfined regions in quasi-periodic quantum wells : DNA nanostructure study

**DOI:** 10.1101/2022.03.25.485884

**Authors:** Alan Tai

**Affiliations:** Science and Life Research Phoenix, AZ85042, USA; Department of Physics, Boston College, Chestnut Hill, MA02167, USA

**Keywords:** quasi-periodic, DNA, quantum well, charge transfer, nanostructure, Fibonacci, Thue–Morse

## Abstract

This study presents a quantum well model using transfer matrix to analyse the charge transfer characteristic in DNA nanostructure sequences. The unconfined state or unbound state above the quantum well are used to investigate the carrier behaviours in nanostructure of semiconductor. Similar analysis and simulations can be applied to the understanding of charge transfer in DNA nanostructure with sequence of periodic and quasi-periodic sequences. The model was validated with experiments using photoreflectance spectroscopy on nanostructure in semiconductor superlattice. In addition, the results published by the study of Thermoelectric effect and its dependence on molecular length and sequence in single DNA molecules[1] were used to compare with the simulations by the quantum well model. The results show agreement and can provide new insight into charge transfer and transport in DNA nanostructure with different types of sequences.

## 1. Introduction

DNA plays an important role for life to develop on earth because it holds the library of information that is critical for the “software” operation within any life form. The charge transfer**[2–4]** within the DNA gives rise to efficient assessments of information and communication for the normal operation of a cell. Similar to a faster CPU and higher transfer rate of data communication inside a computer, the functionality of DNA can be enhanced when the signal speed transmitted along the DNA sequence is increased. Extensive investigation by researchers all over the world have discovered the amazing biochemical mechanisms, charge transfer, and semiconducting characteristics in DNA**[5,6]**. In the microscopic dimension compatible to a quantum well superlattice, the knowledge gained from semiconductor science can be applied and used to supplement to a better understanding of DNA**[7–9]** or vice versa.

Studies of the charge transfer between electron and hole states in multiple quantum well superlattices in semiconductors are topics of great interest in many device applications. Two major factors influencing the signalling rate and charge transfer within the DNA sequences are related to quantum tunnelling in the short range of the under-barrier effect, and the hopping in an extensive range of the above-barrier effect. They relate two major factors influencing the signalling rate and charge transfer within the DNA sequences to quantum tunnelling in the short range of the under-barrier effect, and the hopping in extensive range of the above-barrier effect. The unconfined energy states or unbound states**[10]**-those which lie above the conduction band barriers or below the valence band barriers - play a significant role in carrier capture and in optical transitions.**[11,12]** The quantum effect has influenced even in the region above the barriers. The unconfined energy state has the characteristics of a continuum in the quantum well system, rather than being composed of discrete levels of bound states as in the confined energy region inside the quantum well. However, the quasi-periodic quantum wells in a superlattice structure can give rise to resonance states that have characteristics like those of bound states inside the quantum well.**[13]** The sequence of the quasi-periodic quantum well generated by certain mathematical equations was shown to modulate the unconfined region or above-barrier region so that the energy states have very narrow peaks in the transmission coefficient spectrum. Unlike the periodic quantum well gives rise to broader peaks in the transmission coefficient spectrum, the narrow peaks in the quasi-periodic quantum well show that an electron or hole can become more stable in travelling in the above barrier region. When a carrier entering the unconfined region of a quantum well system with a transmission coefficient equal to unity, the corresponding unconfined energy state adopts a resonance state. Resonant states above the quantum well behave similarly to bound states inside the quantum well and are also called “quasibound” states. The relations for time and energy according to the Uncertainty Principle**[14]** of δtδE~h shows that the discrete-like energy state above the quantum wells can remain in the energy state used for charge transfer for a time longer than those in other energy states with smaller transmission coefficients and broader peaks. Studies of the resonant states in quantum wells and barriers showed that the transmission coefficient pattern with narrow peaks or smaller bandwidths can have physical significance in both superlattice semiconductor and DNA nanostructure.

DNA possesses the curious ability to conduct charge longitudinally through the π-stacked base pairs that reside within the interior of the double helix. In general, the charge transfer is used when a carrier, created (e.g., by oxidation or reduction) or injected at a specific location, moves to a more favourable location, without the application of external voltage. Similar term of charge transport is used when the system is held between electrodes and that a voltage is applied between these electrodes.**[15]** The rate of charge transport through DNA has a shallow distance dependence. DNA is exquisitely sensitive to disruptions, such as environmental change and DNA damage, that affect the dynamics of base pair stacking.**[16]** Perpendicular to each pairing are adjacent nucleotides whose heterocycles’ π-orbitals overlap, creating a path for charge to transport down the helical axis.

In this work, I studied the charge transfer in periodic, Fibonacci and Anti-symmetric DNA sequencies by applying the quantum well model of transfer matrix techniques. This quantum well model was validated by comparing the results published by the study of Thermoelectric effect and its dependence on molecular length and sequence in single DNA molecules.**[1]** In addition, the experimental techniques of photoreflectance spectroscopy (appendix) were used to investigate the optical transitions between the unconfined states in different superlattice systems.**[13]** The spectra reveal these transitions are significantly enhanced in Fibonacci superlattices and some quasi-periodic superlattices over periodic and random systems of a similar composition. The experimental results are compared with transition energies computed by the quantum well transfer matrix technique to get the transmission coefficients in different energy states. When the transmission coefficients peaks are narrow and approaching to 1, the charge transfer of the carriers are also increased. The same computational method was used to calculate the functionality of the charge transfer in the DNA systems.

This paper is organized as follows. Kronig-Penny model and transfer matrix method of the quantum wells in DNA sequences are presented in §2. DNA with periodic sequences are simulated to compare with the charge transfer characteristic of quasi-periodic sequences in §3. The validation of the model by the study of Thermoelectric effect in DNA sequences, Bloch wave vector and photoreflectance measurements on different quantum well superlattices (appendix) are presented in §4. Discussion and proposal of future works in both computer simulation and experiment measurements for charge transfer and transport in DNA are presented in §5. The conclusion of this study is presented in §6. The last session in the appendix reviews some of the previous work for the experimental techniques of photoreflectance spectroscopy that were used to investigate the optical transitions between the unconfined states in different superlattice systems.

## 2. Quantum Wells modelling of DNA

Kronig and Penny used a one-dimensional square well periodic potential to calculate the energy band of a general periodic bulk crystal. The same principle can be applied to the quantum well system with the periodic sequences and also extended to the non-periodic sequence using the effective mass approximation and the transfer matrix technique. The transfer matrix technique takes advantage of the convenience of applying computer methods to the solution of matrixes numerically. The transmission coefficient can then be found to determine the unconfined electron subband energies in the above barrier region and confined electron subband energies within the quantum wells.

Referring to figure 1, the positions of the barriers and wells are located at xm and xn respectively in the x-axis, where the series 1,2,3,4…,m, n,… are integers in ascending order. The effective mass m* in region M(well) and region N(barrier) are mw and mb respectively, where the series 1,2,3,4…M,N,… are integers in ascending order. The solutions of the Schrodinger equations in the quantum wells are given as:

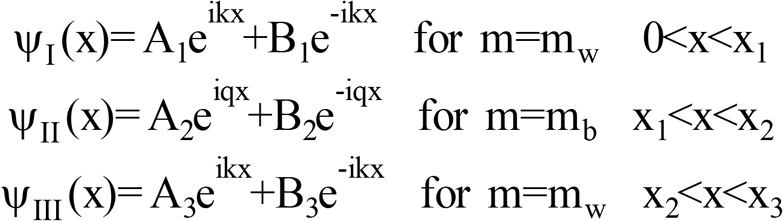

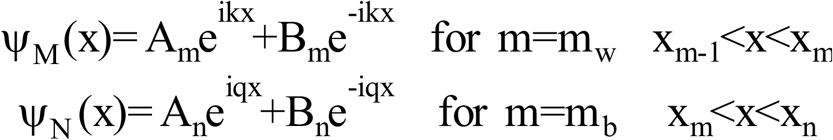

**Figure 1.**
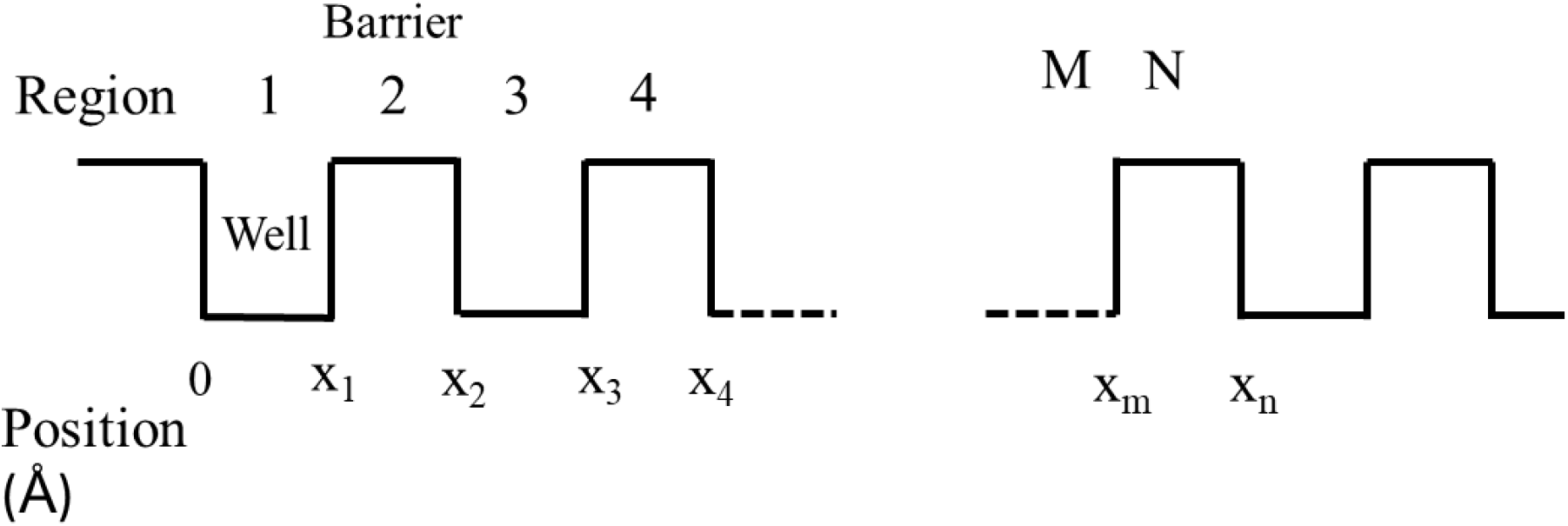
The DNA nanostructure sequence along the x-axis.

The solutions of the Schrodinger equations in the various regions are

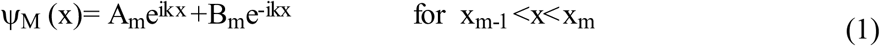

where 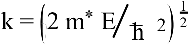 corresponding to the wave number in the well region.

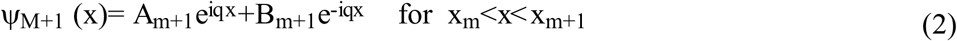

where 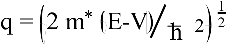 corresponding to the wave number in the barrier region, E is the energy above the bottom of the well and V is the potential at the top of the barrier. The energy above the bottom of the well is designated, E, and V is the potential at the top of the barrier. By employing the boundary condition for continuity of the wavefunctions and their derivatives, we can form a sequence of 2 x 2 transfer matrices.

The transfer matrix corresponding to the boundary located at x_m_ is:

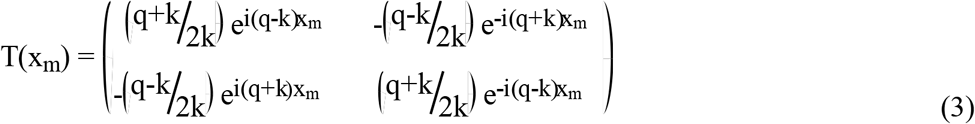

The transfer matrix corresponding to the boundary located at x_m+1_ is:

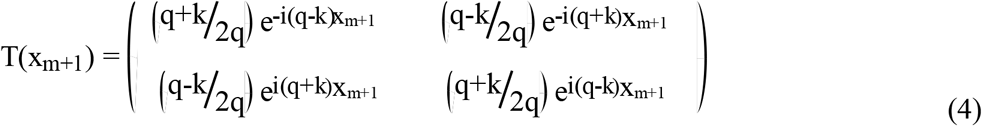

The final transfer matrix is given by the product,

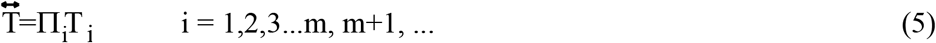

and the corresponding transmission coefficient, t = 1/T_11_, where T_11_ is the final diagonal component of the matrix 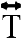.

## 3. Simulation results

We simulated Four different sequences based on the 4 types DNA’s nucleotides, adenine (A), thymine (T), guanine (G), and cytosine (C) for the unconfined states in above barrier region using the transfer matrix techniques described in §2. Periodic sequenced DNA was first simulated and then two non-periodic sequenced, or quasiperiodic DNA: Fibonacci sequence, Anti-symmetric sequence are simulated.

In order to build relations between the charge transfer of DNA and simulation model, we used one-dimensional sequence with 1 or 0 to simulate the sequence of quantum wells and barriers respectively for the corresponding nucleotides. The DNA sequences under study are considered composing of repeated stacks of nucleobases formed by either G-C/C-G as the well (1) or A-T/T-A as the barrier (0). The energy levels between G-C and A-T base pairs are shifted by about 0.4 eV**[17]**. The difference in energy level forms the quantum wells and barriers, as in figure 2. Both of the base pairs have the same nucleotide length of 3.4Å for each basic unit of 0 or 1 that counted as 1 layer in the simulation parameter. The helical chains of nucleotides in DNA are bounded to each other by hydrogen bonds that coil into tight loops and to form different shapes of polymers. Conductivity has been found to be dependent on sequence, hydration, length, temperature, and hybridization in some experiments. The environmental and helical factors are not considered and are assumed to be constant among different DNA sequencies for comparison under this basic 1-D model. Table 1 shows the effective masses of electrons and holes along directions perpendicular to the staking planes (in unit of the free electron mass m_0_)[18]. The effective masses of electrons **m**_**e**_ of 5 is used for the simulation of charge transfer based on the average of G-C combinations in table 1. Note that the **m**_**e**_ and **m**_**h**_ have the same order of magnitude for the DNA nucleotides. The **m**_**e**_ is an empirical value determined by experiments and the calculated lattice parameters. This value can be adjusted for environmental factors change to match the simulation results and experiments, as discussed in a future work of §5.

**Table 1.**
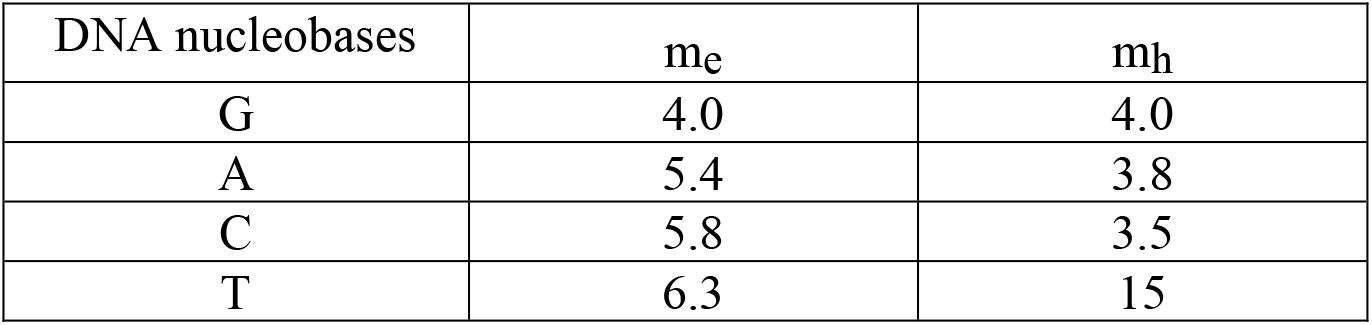
Anhydrous crystals of DNA nucleobases with effective masses of electrons and holes along directions perpendicular to the stacking planes (in units of the free electron mass m_0_) are shown.**[18]**

**Figure 2.**
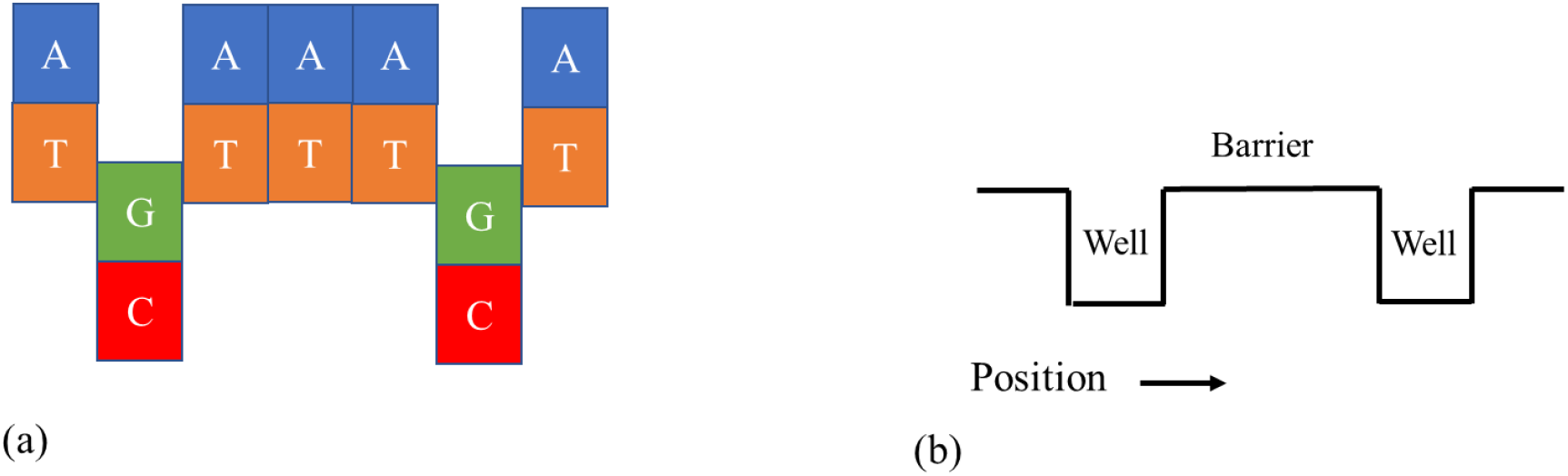
Schematic representation of the energy profile in a DNA with 7 base pairs. (a) Ionization potential for G-C base pairs is differ from A-T base pairs of about 0.4 eV. (b) System (a) can be treated as a series of quantum wells and barriers systems in different positions.

### 3.1. Periodic DNA

We first simulated the periodic sequence DNA, with alternating layers of 0 and 1, for the energy states above the quantum well as in the figure 3. Periodic structure is periodically arranged like 011011011011011011011… 44 boundaries and 172 boundaries are selected to compare with other types of DNA using similar numbers of layers. Each layer of nucleotide is corresponding to either 1 or 0 according to the periodic sequence. Each barrier (0) or well (11, well width of 3.4Å x 2 = 6.8Å) has two boundaries to the left and to the right side, as shown in the figure 3. We used an analytical approach described in §2 to formulate for this simulation.

**Figure 3.**
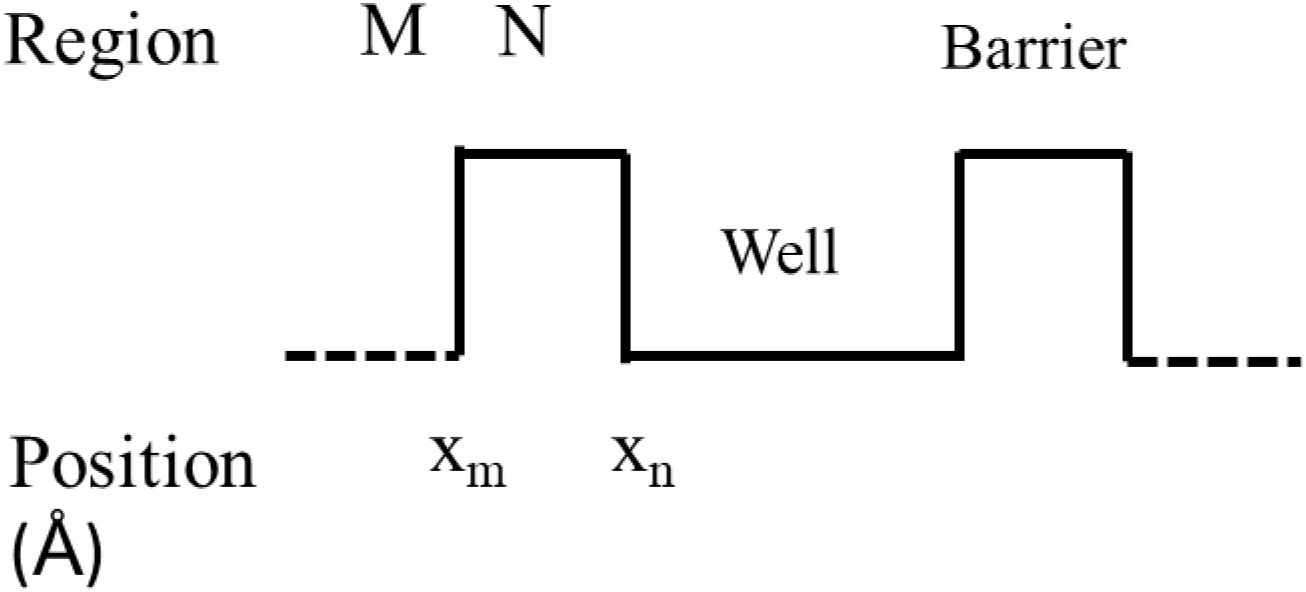
The periodic DNA nanostructure sequence with repeated 011011011… along the x-axis to form different boundaries of DNA.

The simulation results in figure 4a and figure 4b show the transmission coefficient vs energy level for 44 boundaries and 172 boundaries respectively, in meV above the bottom of the QW in the conduction band structure of the DNA sequence. Our focus is on the region above the barrier height of 400meV that there is a gap from 400meV to 515meV, followed by a band up to 764.1meV. There are subsequent gaps and bands after the first band depending on how far into the continuum of the unconfined region are simulated. The number of peaks oscillating in the band increase with the number of boundaries increase from 44 boundaries to 172 boundaries as calculated by the computer simulation. However, the corresponding positions of gaps and bands remain the same when compared between the two simulation results in figure 4(a) and figure 4(b).

**Figure 4.**
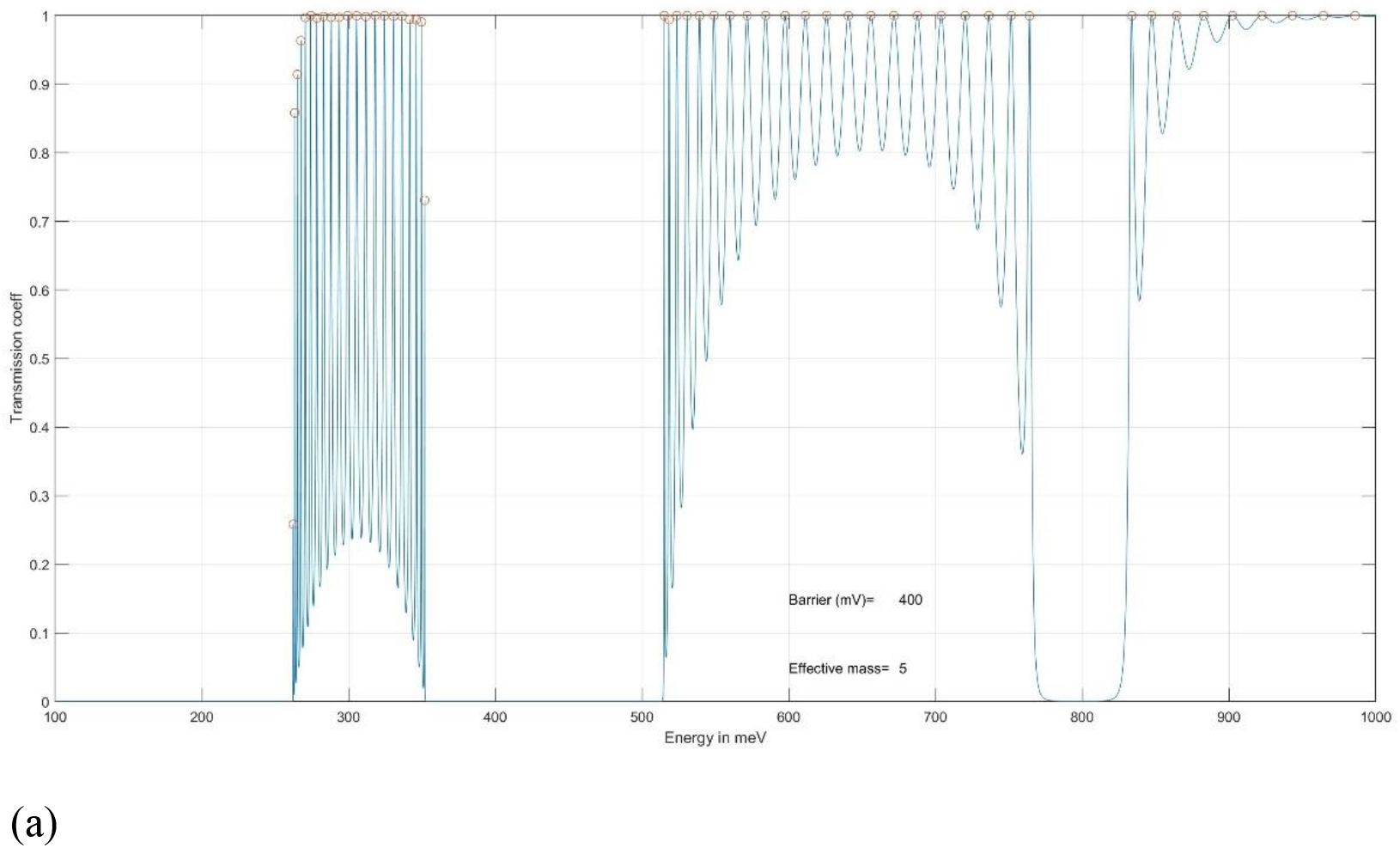
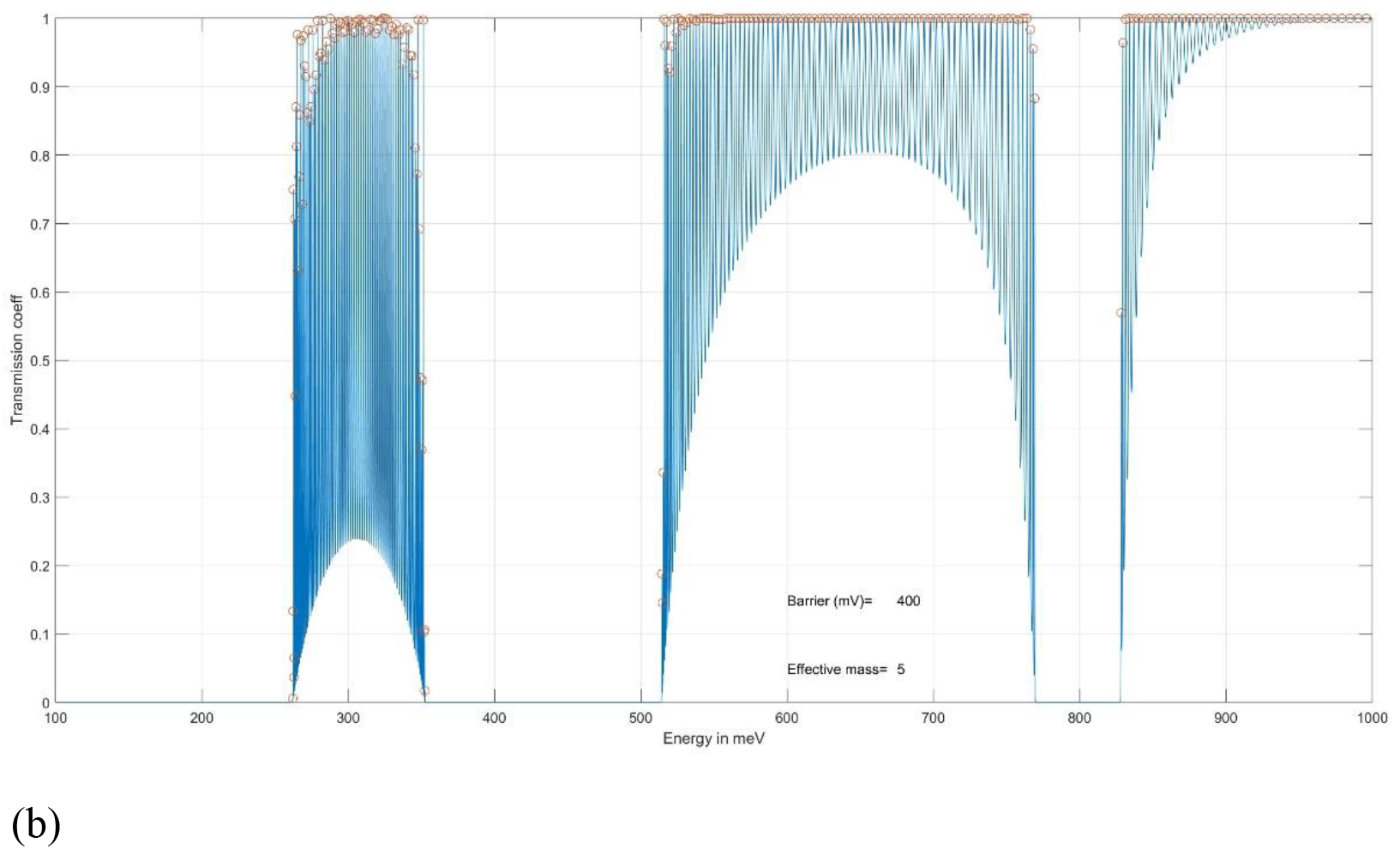
Computer simulation of Transmission coeff. Vs Energy for periodic DNA, quantum barrier = 400mV and electron effective mass **m**_**e**_ = 5 with (a) 44 boundaries (b) 172 boundaries.

### 3.2. Fibonacci DNA

Quasiperiodic sequence DNA, with layers arranged according to the Fibonacci sequence, is shown in table 2. The Fibonacci sequenced DNA comprises an arrangement of layers of 0 and 1 following the Fibonacci sequence: S1 = 0, S2 = 1, S3 = 01, S4 = 101,…, Si = (Si-2) (Si-1), Si+1 = (Si-1) (Si). This leads to a quasi-periodic sequence of wells and barriers with Fibonacci structure formed into a Fibonacci arrangement of 01011010110110101101011… where the third group of sequences was formed by the two former groups of sequences combined. In this Fibonacci sequence of 0 and 1, there are wells with only one layer of “1” and maximum of two layers of “1” together to form the smaller quantum well and larger quantum well respectively. There are two boundaries for each of the large and small quantum well but have two layers of nucleotides in the large quantum well and only one layer of nucleotide in the small quantum well. There is only one layer of “0” with two boundaries in between the quantum wells of the Fibonacci DNA. In addition, there is mirror image at layer 28 with layer 1 – 27 vs layer 29 – 55, which corresponds to the 44th boundaries for the quantum well and barrier sequence.

**Table 2.**
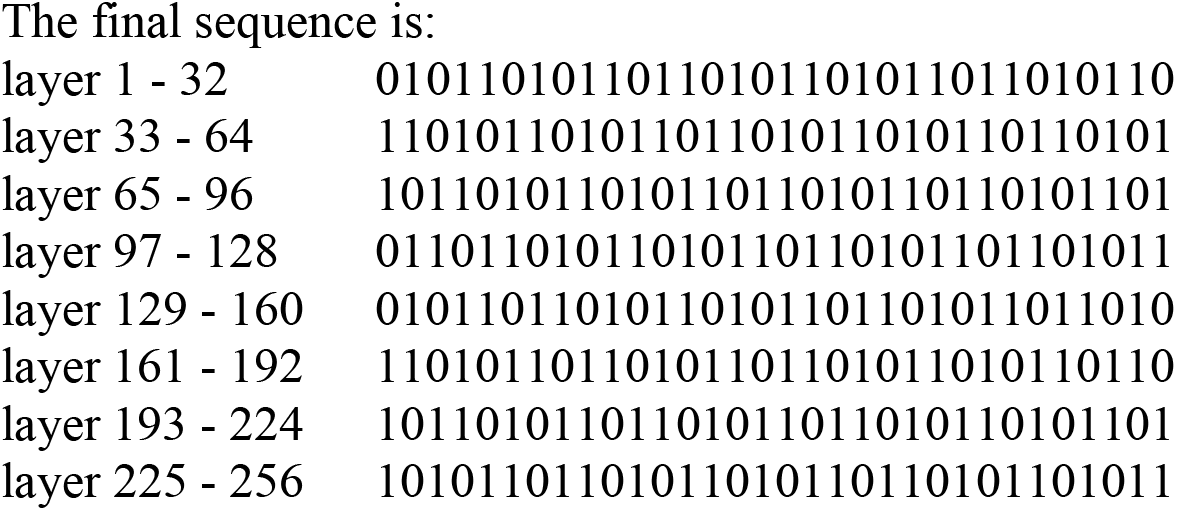
Fibonacci sequenced DNA

The simulation results in figure 5 (a – c) show the transmission coefficient vs energy level for (a) **m**_**e**_ = 5 with 44 boundaries (b) **m**_**e**_ = 5 with 100 boundaries (c) **m**_**e**_ = 3.5 with 44 boundaries respectively, in meV above the bottom of the QW in the conduction band structure of the Fibonacci DNA sequence. There are distinct and narrow transmission peaks separated from each other inside the quantum well region (<400meV) and in the initial band in the unconfined region (>400meV). When the simulation boundaries increase from (a) 44 to (b) 100 for **m**_**e**_ = 5, the transmission peaks inside the quantum well reduce, but the transmission peaks increase in the unconfined region. The corresponding positions of gaps and bands remain the same when compared between the two simulation results in figure 5(a) and figure 5(b). In figure 5(c), **m**_**e**_ = 3.5 with 44 boundaries is used for simulation to see how the change in **m**_**e**_ affects the transmission peaks. The simulation result shows that the relative position of the band and gaps are shifted to the right. There are also some transmission peaks appear just above the quantum well into the unconfined region.

**Figure 5.**
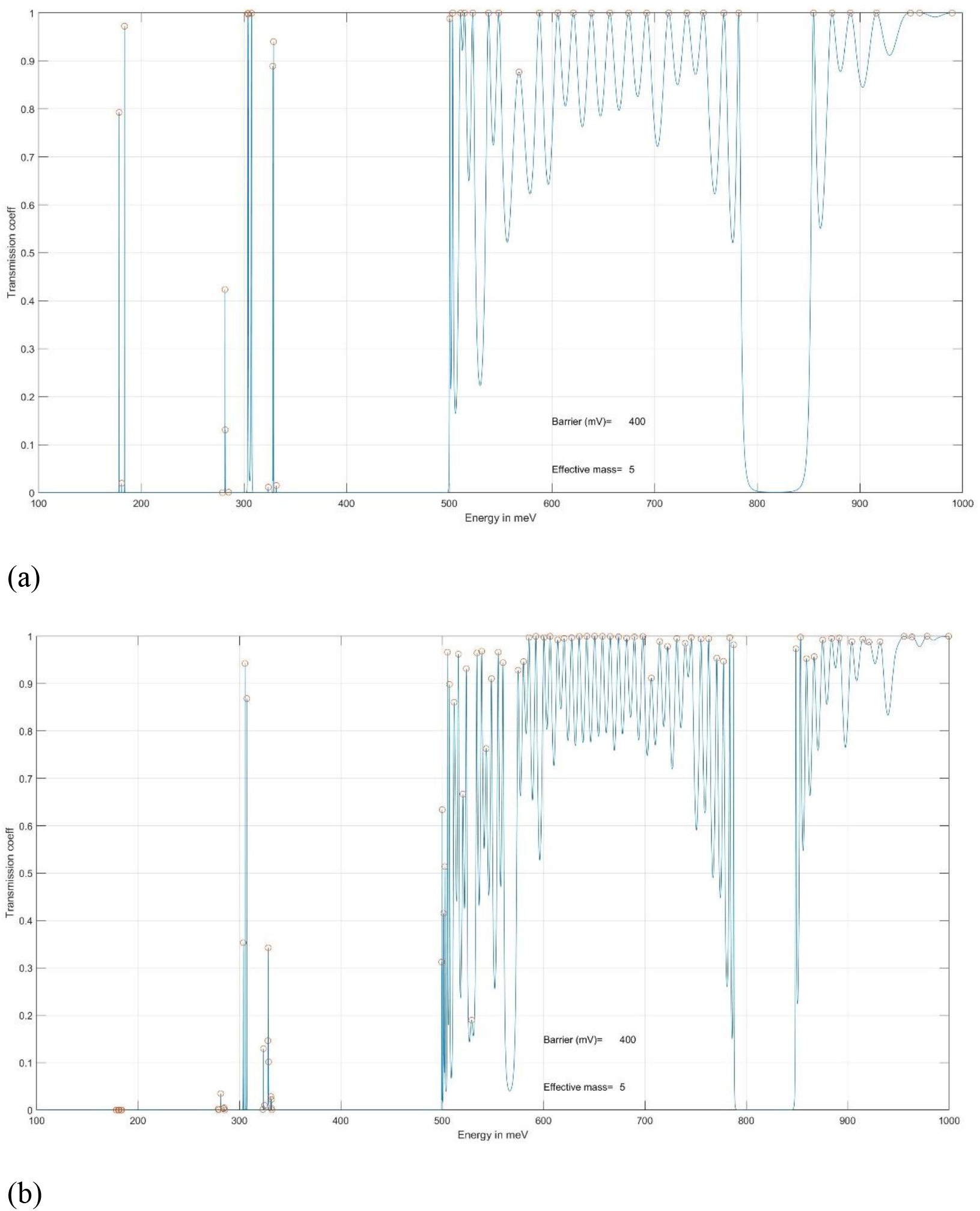
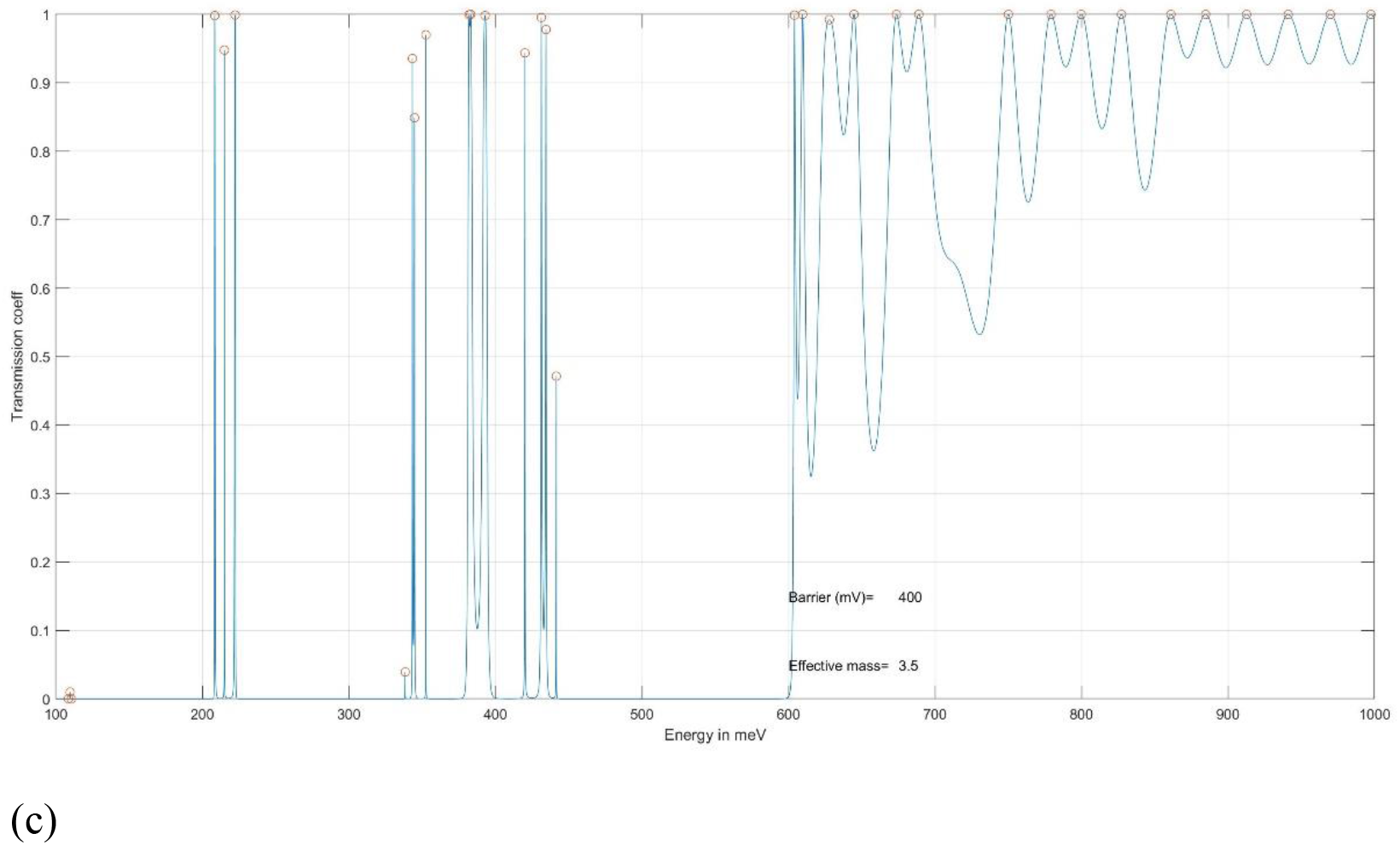
Computer simulation of Transmission coeff. vs Energy for Fibonacci DNA, quantum barrier = 400mV and (a) **m**_**e**_ = 5 with 44 boundaries (b) **m**_**e**_ = 5 with 100 boundaries (c) **m**_**e**_ = 3.5 with 44 boundaries.

### 3.3. Anti-symmetric DNA

Anti-symmetric sequenced (also called Thue–Morse**[19]** sequence) DNA, with layers of barriers and wells arranged according to a special sequence which “Anti-Symmetric” sequence, is shown in table 3. The basic idea is to form a sequence which is opposite (barrier becomes well; well becomes barrier) to the entire sequence before it.

For example:

First layer is 0 (barrier), second layer is 1 (well),

third and fourth layer will be 10(well and barrier),

fifth to eighth layer will be 1001(well, barrier, barrier and well).

Therefore, the sequence will be:

0 1 10 1001 and so on

**Table 3.** Anti-symmetric sequenced DNA

|  |  |
| --- | --- |
| layer 1-32 | 01101001100101101001011001101001 |
| layer 33-64 | 10010110011010010110100110010110 |
| layer 65-96 | 10010110011010010110100110010110 |
| layer 97-128 | 01101001100101101001011001101001 |
| layer 129-160 | 10010110011010010110100110010110 |
| layer 161-192 | 01101001100101101001011001101001 |
| layer 193-224 | 01101001100101101001011001101001 |
| layer 225-256 | 10010110011010010110100110010110 |

Arrangement of layers 1 - 64 is the same as layers 193 - 256; layers 65 - 128 is the same as layers 129 - 192.

The simulation results in figure 6a and 6b show the transmission coefficient vs energy level for (a) **m**_**e**_ = 5 with 44 boundaries and (b) **m**_**e**_ = 5 with 172 boundaries respectively, in meV above the bottom of the QW in the conduction band structure of the Anti-symmetric DNA sequence. There are distinct and narrow transmission peaks separated from each other inside the quantum well region (<400meV) and in the initial band in the unconfined region (>400meV). When the simulation boundaries increase from (a) 44 to (b) 172 for **m**_**e**_ = 5, the transmission peaks inside the quantum well stay the same position, but the number of transmission peaks increase (oscillation) in the unconfined region. The corresponding positions of gaps and bands remain the same when compared between the two simulation results in figure 6(a) and figure 6(b).

**Figure 6.**
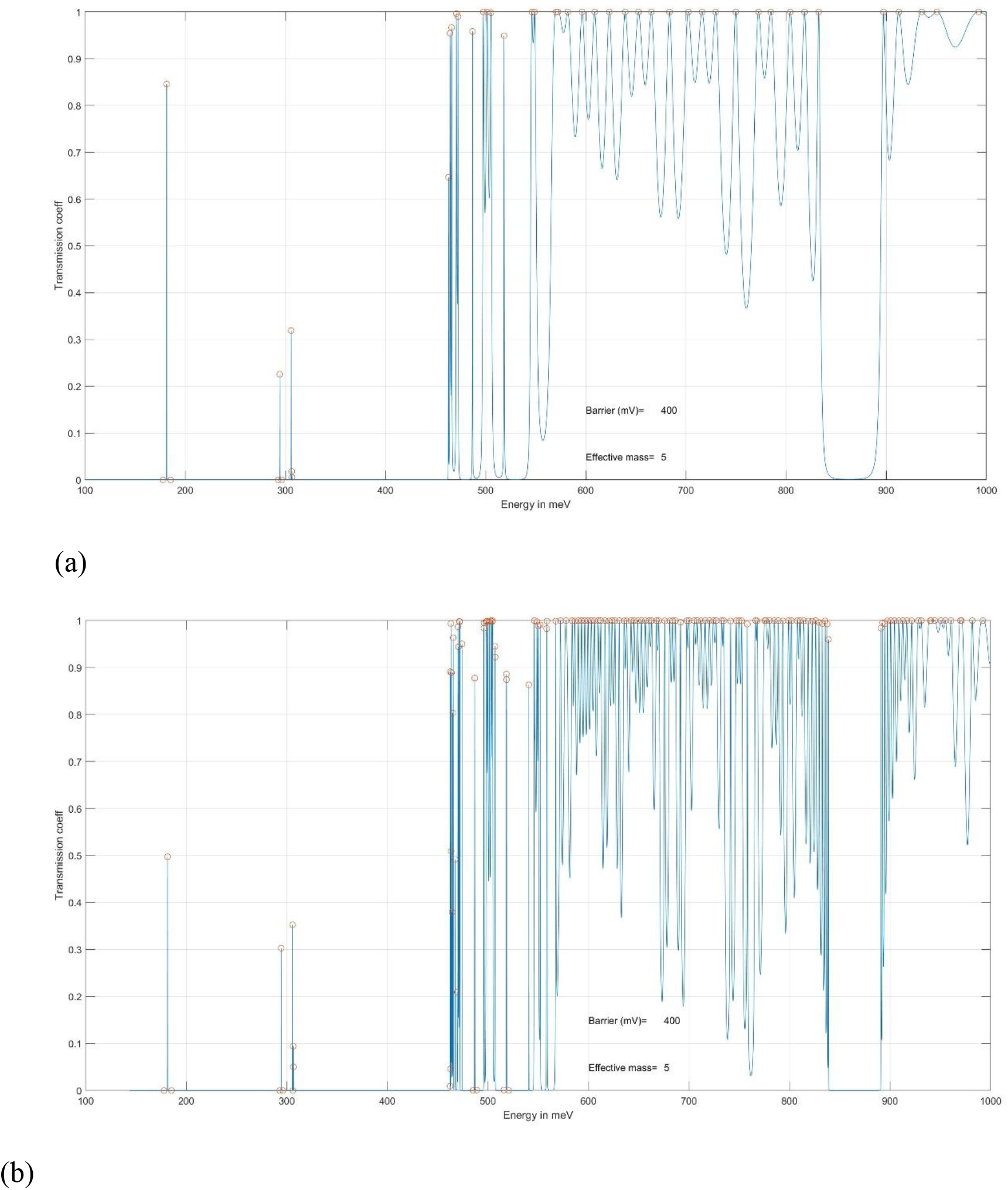
Computer simulation of Transmission coeff. Vs Energy for Anti-symmetric DNA, quantum barrier = 400mV and electron effective mass = 5 with (a) 44 boundaries (b) 172 boundaries.

## 4. Validation of the model

The quantum well model is first used to determine if the simulation of the DNA sequences can show agreement with the related experimental results. Even though the quantum effect on the nanostructure of superlattices is quite similar to that on the DNA structure, there are many factors, such as the chemical composition and environment that can generate different simulation results. Therefore, I will focus this study on the intrinsic effects on the DNA by comparing different DNA sequences while all other key parameters will be treated as constant. All the simulations or experiments will be performed under the same environmental and thermal conditions with the variation factor being in the different DNA sequences.

The results published by Li et al.**[1]** are used to compare with the simulations obtained from the quantum well model of this study. The Seebeck coefficients of DNA in the hopping regime (in A(CG)nT) are small, and weakly depend on the molecular length compared to other organic molecules. By inserting a short AT block (shorter than 5 AT base pairs) into the middle of A(CG)nT, it leads to a much greater Seebeck coefficient, which increases with the AT block length. However, when the AT block is longer than 5 AT base pairs, the Seebeck coefficient drops to the level of A(CG)nT, and become insensitive to the AT length. This transition coincides with the tunneling-hopping transition near 4-5 AT base pairs observed from the conductance measurement, which strongly suggests that the thermoelectric effect is large in the tunneling regime and small in the hopping regime.

Figure 7 (a - d) shows the computer simulation of Transmission coefficient vs Energy for different DNA length (a) A(CG)_3_T (b) A(CG)_5_T (c) A(CG)_6_T (d) A(CG)_7_T, quantum barrier = 400mV and electron effective mass = 5. All the plots have distinct peaks in both quantum wells region (<400meV) and unconfined region (>400meV). The transmission peaks inside the quantum wells correspond to the tunneling regime based on the theory of quantum mechanics. The transmission peaks outside the quantum wells in the unconfined region correspond to the hopping regime. These results show only qualitatively that there are combinations of both tunneling and hopping transitions in the DNA sequences.

**Figure 7.**
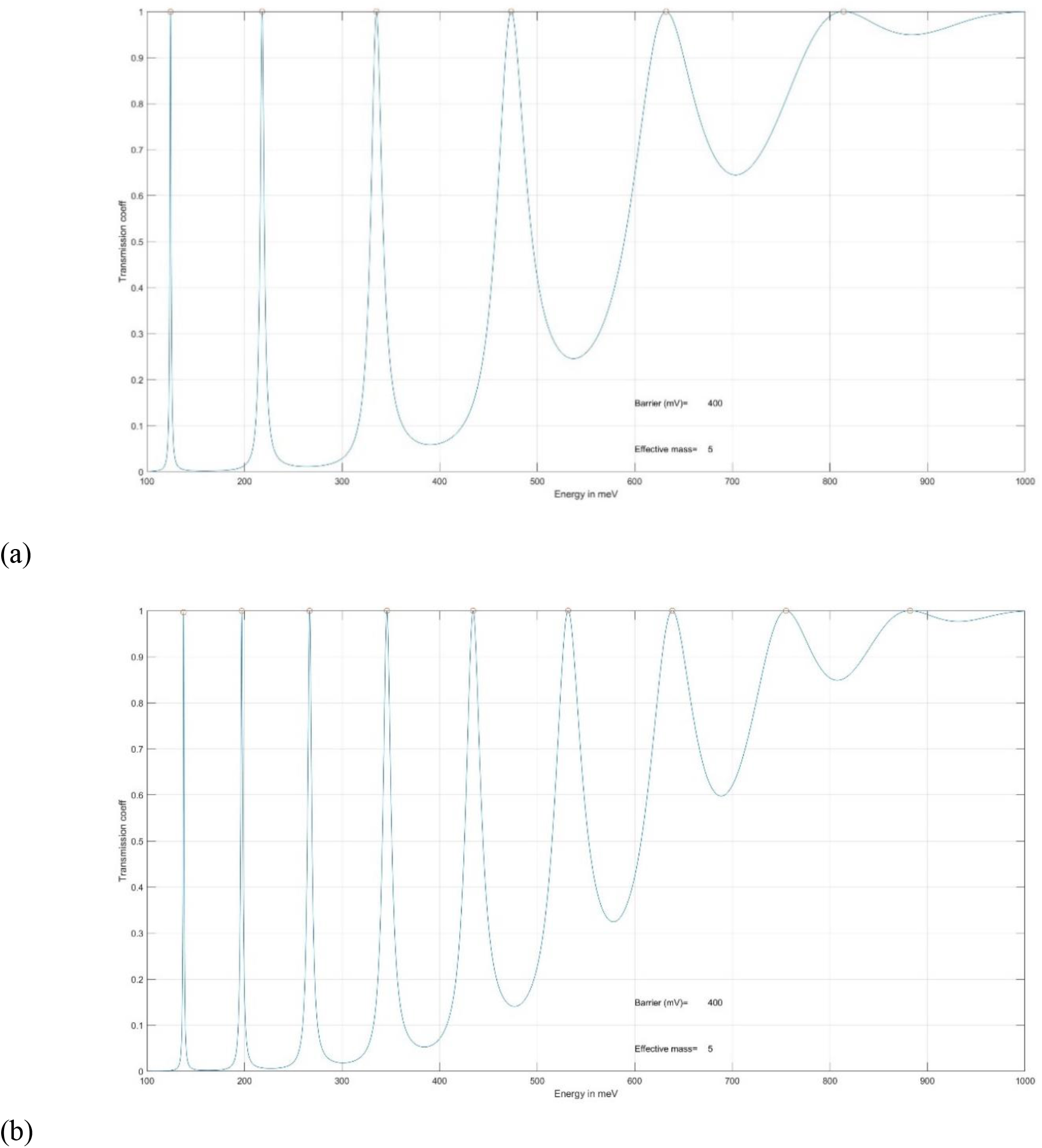
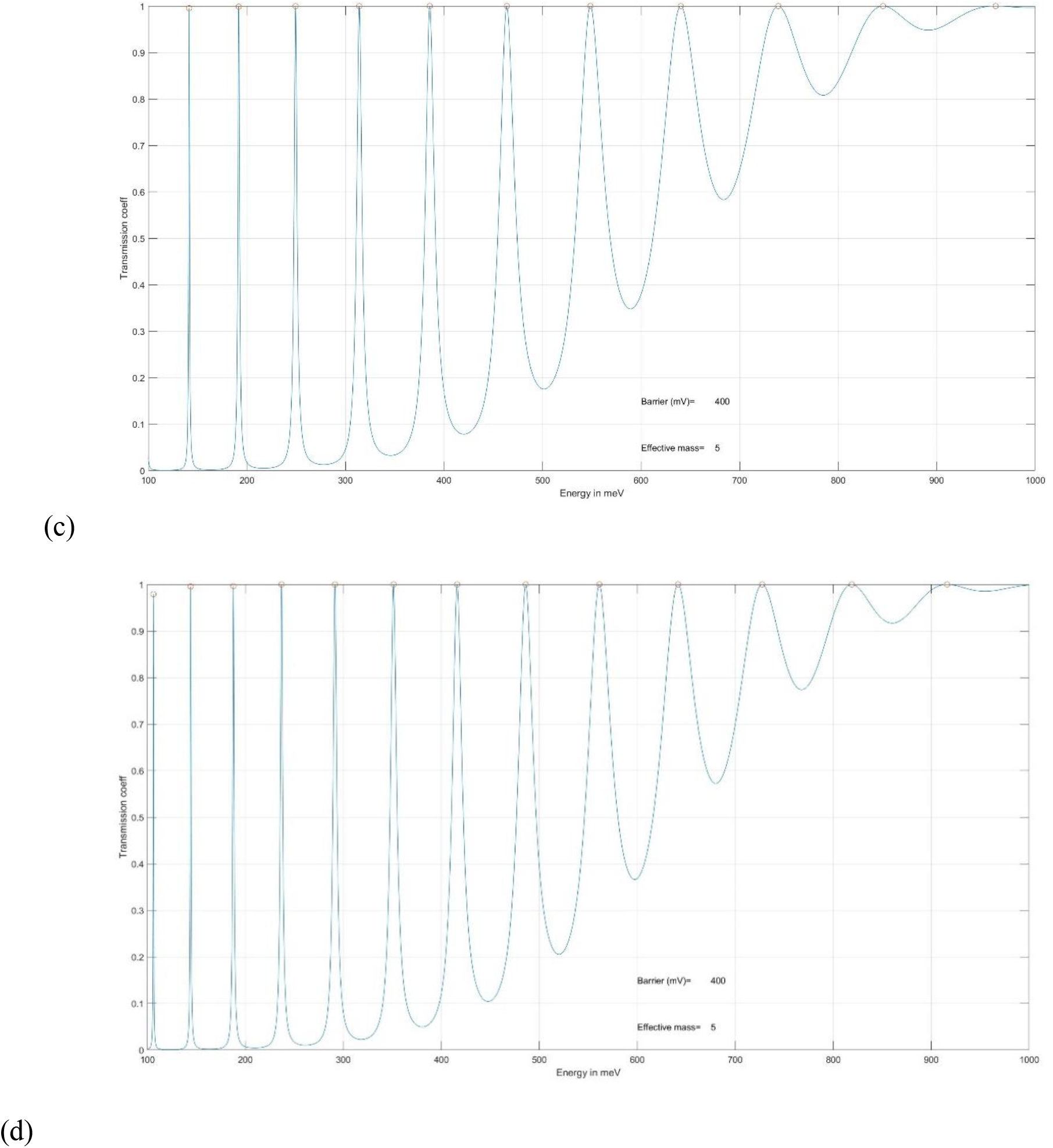
Computer simulation of Transmission coeff. Vs Energy for different DNA length (a) A(CG)_3_T (b) A(CG)_5_T (c) A(CG)_6_T (d) A(CG)_7_T, quantum barrier = 400mV and electron effective mass = 5.

Figure 8. (a – g) shows the computer simulation of Transmission coefficient vs Energy for different DNA lengths in the following: (a) ACGCAGCGT (b) ACGCATGCGT (c) ACGCATAGCGT (d) ACGC(AT)_2_GCGT (e) ACGC(AT)_2_AGCGT (f) ACGC(AT)_3_GCGT (g) ACGC(AT)_4_GCGT quantum barrier = 400mV and electron effective mass = 5. By inserting a short AT block (a – d, shorter than 5 AT base pairs) into the middle of A(CG)nT, the transmission peaks inside the quantum wells (E <400meV) become progressively reduced to a smaller peak in figure 8d (4 AT base pairs). With 5, 6 and 8 AT base pairs, the transmission peaks inside the quantum wells(E <400 meV) gradually disappear (t < 0.08) but the unconfined region(E >400meV) retains some well-defined transmission peaks (t = 1) as showed in figure 8 (e – g).

**Figure 8.**
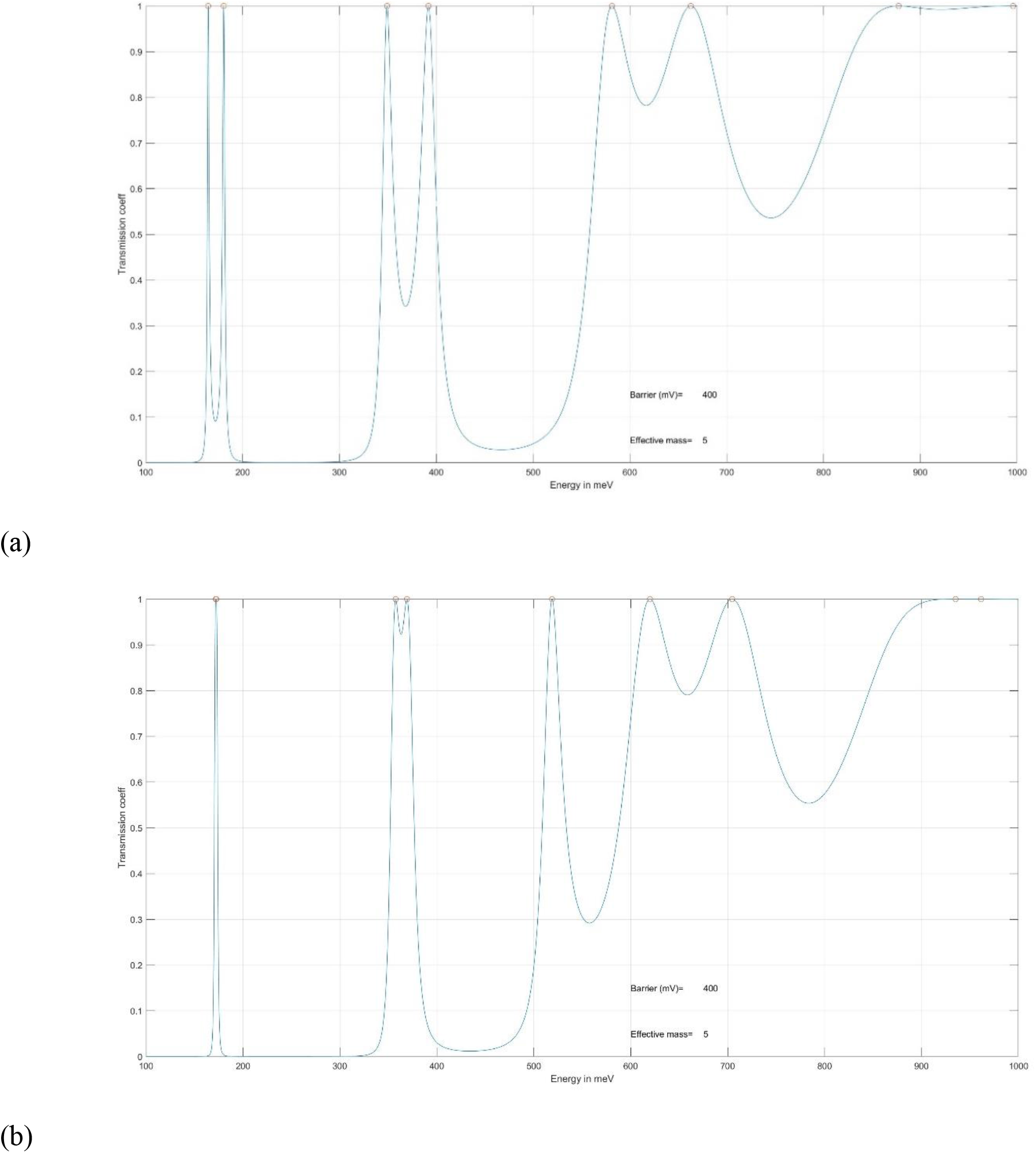
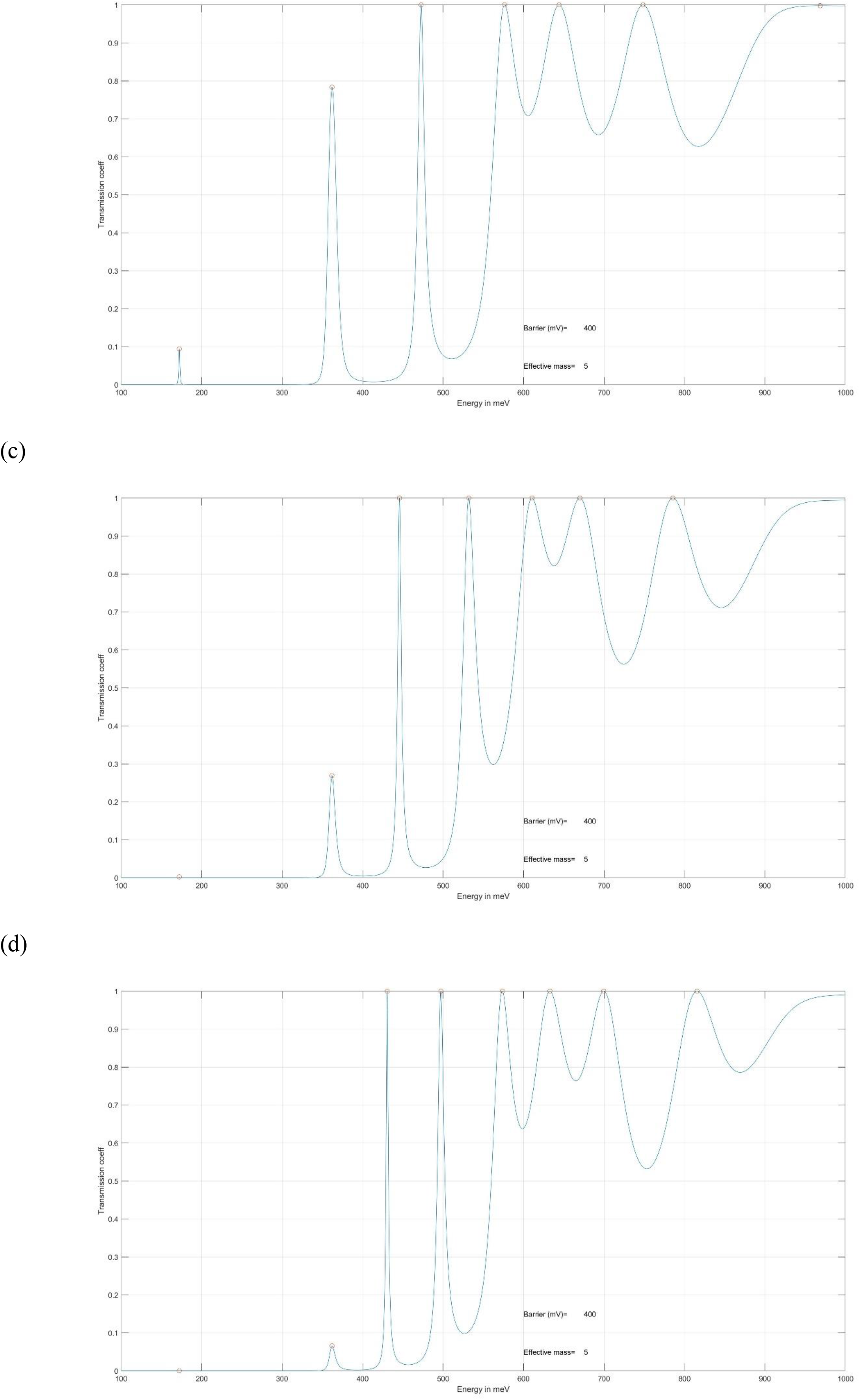
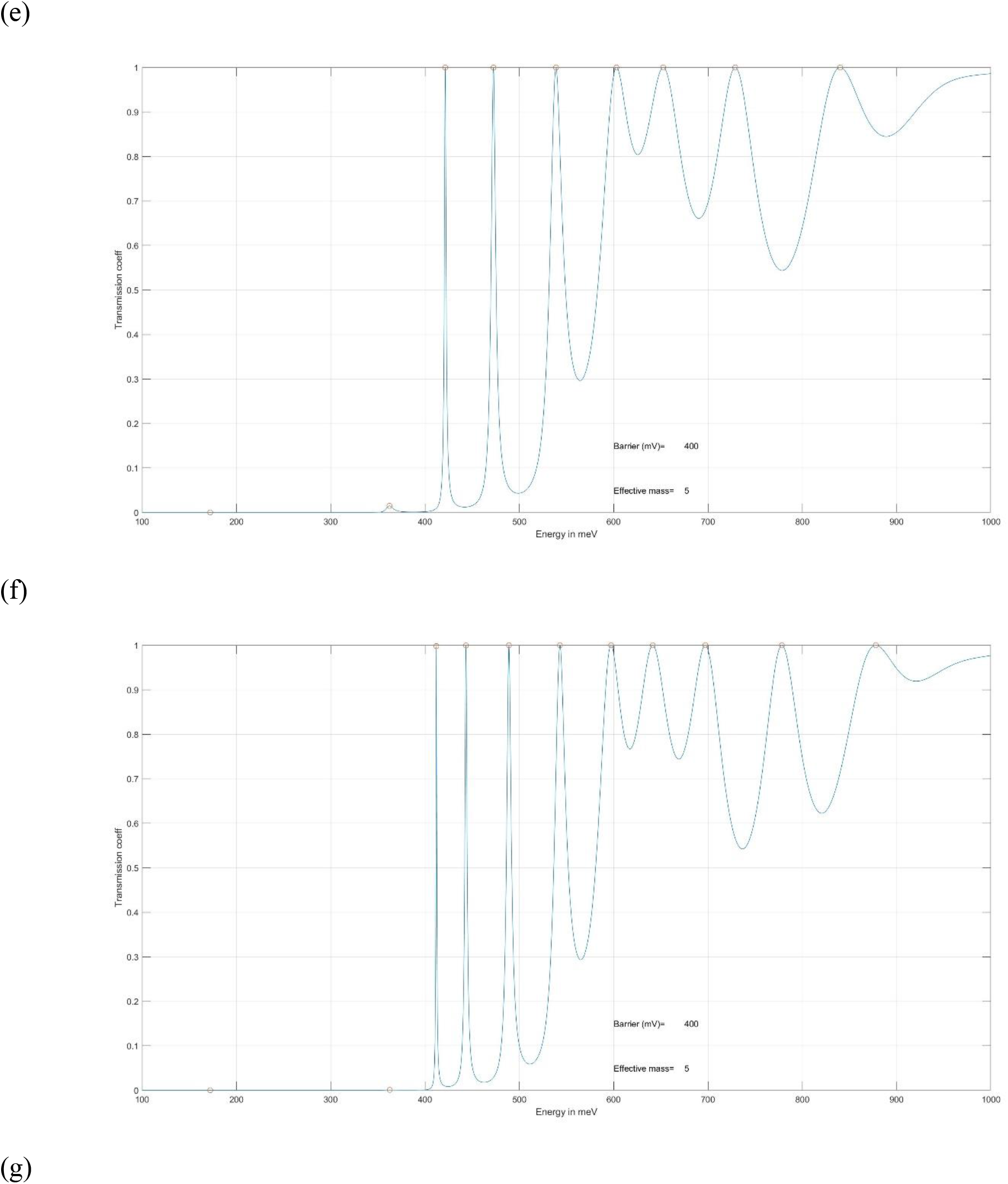
Computer simulation of Transmission coeff. Vs Energy for different DNA length (a) ACGCAGCGT (b) ACGCATGCGT (c) ACGCATAGCGT (d) ACGC(AT)_2_GCGT (e) ACGC(AT)_2_AGCGT (f) ACGC(AT)_3_GCGT (g) ACGC(AT)_4_GCGT quantum barrier = 400mV and electron effective mass = 5.

The computation results of the quantum well model for the different DNA length samples show that the prediction appears to be valid in the tunneling regime and in the hopping regime (unconfined region). For DNA with the inserted A, AT, ATA or ATAT as barrier in the middle of A(CG)3T, the experimental results of large Seeback coefficient (S: 5 to 7.9 μVK^−1^) agree with this quantum well model of transmission peaks regarding the tunneling effect within the quantum well regions formed by the barriers. For DNA with the inserted AT base pairs > 4 as barrier in the middle of A(CG)3T, the experimental results of smaller Seeback coefficient (S: 4.9 to 2 μVK^−1^) agree with this quantum well model of transmission peaks regarding the hopping transport above barriers. Figure 8f and 8g show there are only narrow and transmission peaks with t =1 appear above the barrier of E >400meV while there have no transmission peaks for E < 400meV. Good agreement between the experiments[1] and the simulation of varying the AT block combinations for the charge transfer of both tunneling and hopping of the DNA sequences is demonstrated. This agreement and application will be further discussed in §5.

## 5. Discussion and proposal for future work

The investigation of the charge transfer and transport mechanisms in molecular DNA structures and those of nanoelectronic devices can lead to a greater understanding of the communication processes in the two. However, the environmental and chemical factors give rise to many challenges that affect the carrier motions in the communication for short and long range charge transport. In order to minimise the known factors affecting the physical sequences, I propose Quantum Well analysis be applied on the unconfined region to the nanostructure for both the periodic and quasi-periodic system. The Quantum Well semiconductor superlattice is typically in the scale of nm, about 10 times the DNA unit where a similar Quantum Mechanics can be found. By focusing on the quasi-periodic sequence and comparing with the periodic and random sequence of Quantum Well structures in superlattices, the unconfined states are verified through experiments (see Appendix) to enhance charge transfer and transport**[13]**. Quantum Well Transfer Matrix Method (TMM) was used to demonstrate that the basic model facilitates the computational process of the transmission coefficient and the energy positions of transitions. Similar approaches can be used to compare and model different sequences of DNA. The computer simulation presented here is just a starting point for providing more experiments to calibrate and refine the model, such as the variation of the effective mass of the carrier. This is assuming the environment and chemical factors that change the CT can be put in a stable constant situation so that relative values related to the outcomes among different types of sequences can be compared.

Quantum wells in a superlattice can simplify the investigation in a similar sequence in DNA, which has more variance in factors that can influence the outcomes, cross check and optimize first to complement result of using DNA molecules. Comparing with the DNA sequences, the quasi-periodic sequence of non-periodic and non-random arrangement can be generated in a semiconductor superlattice to investigate if certain DNA sequences can impact characteristic of charge transfer. There are many factors (e.g., aqueousness, counterions, extraction process, electrodes, purity, substrate, structural fluctuations, geometry) influence the carrier motion along DNA. These factors are either intrinsic or extrinsic. Here, the focus is on the most important of the intrinsic factors, i.e., the effect of alternating the base-pair sequence, which affects the overlaps across the *π*-stack. The aim of this work is a comparative examination of the influence of base-pair sequence on charge transfer, in aperiodic or quasi-periodic sequences.**[15]**

Using the theoretical analysis for the unconfined region of the superlattices, I apply a similar investigational model to simulate the transmission coefficient spectrum for the unconfined region in the DNA sequences. The analysis of which can be verified through experiments presented in this paper. The presence and absence of water have been shown to influence on the band structure of the DNA base stacks. However, the goal of computational studies is mainly used to understand the impact of different DNA sequences on the charge transfer within the DNA. The simulation results of the DNA sequences that provide enhanced functions of charge transfer are proposed to be used for future experiments in charge transfer. The verification and further studies of different aperiodic DNA sequences will provide more insights into the charge transfer of DNA and will optimize the electronic transfer rate regarding the energy states in the above-barrier of the QW unconfined regions.

Based on the agreement between this quantum well model and the experimental studies by Li et al.[1], my hypothesis is that the transmission peak simulation is an effective tool to supplement the analysis by hopping mechanism and other transport theories**[20]** in DNA sequences. The application of TMM not only can be used for few DNA base pairs but also can be implemented for thousands or more base pairs like those of the human gene. The computation time is usually less than 1 minute depending on the types of computers are used. The complexity of the Quantum Well model can be expanded to include the extrinsic factors and the computation time would be increased. The transmission peaks and their bandwidths in the transmission coefficient simulation can correlate with the tunneling/hopping resistance and thermoelectric effect in the DNA molecules. Below steps can be used to obtain the optimised DNA sequences that cause the CT between the donor and acceptor of carrier to proceed through the fastest pathway available in the unconfined region for longer range DNA CT.

1. Using the Quantum Well model presented here for simulation to get the transmission coefficient vs energy for different quasi-periodic and random sequences of Quantum Well nanostructures.
2. Picking the narrower bandwidths with transmission peak =1 as selection criteria for optimum sequences in DNA and superlattices.
3. Both of the superlattices and DNA in step 2 can be built for the below samples with emphasis on step 3b or in case resources of step 3a are not available.
  a. Building the quantum well superlattice (see examples in the appendix) according to step 2 and using the Photoreflectance or other techniques to find the Quantum Well sequences that has best CT characteristic.
  b. Prepare the DNA molecules (see example of the studies by Li et al.) according to step 2 and using the STM, conductivity measurement or other techniques to find the Quantum Well sequences that has best CT characteristic. The hopping resistance and Seeback coefficient can be measured to determine which sequences of the DNA would have the smallest values or the thermoelectric effect vanish.
4. Studying above steps systematically and using the experimental results as feedback to optimize the Quantum Well model and simulation based on the empirical values of the effective mass, barrier height and other related parameters.

The advantages of the above studies are to apply the verified Quantum Well model for intrinsic characteristics of the DNA molecules with different sequences and lengths that can result in effective approaches of searching for the sequences optimised for CT in nanostructures. The examples in the appendix demonstrate the quantum well sequencing has special quantum mechanics influence on the unconfined states. I have searched for hundreds to thousand of different combinations of sequences by simulation in order to pick the Anti-symmetric and Fibonacci sequences as the appropriate sequences to be built into actual superlattices**[13]**. These quasi-periodic sequences have unique mirror like symmetric at certain particular location and may well be suitable to facilitate the CT in DNA molecules. The double stranded structure in DNA run in opposite directions to each other and is thus antiparallel. The studies of Anti-symmetric sequences of DNA with antiparallel characteristic can provide more insight into the CT behaviours.

Another interesting studies**[21]** by Xiang et al. of applying an electrochemical (EC) gate voltage to the DNA molecule, leading to reversible switching of the DNA conductance between two discrete levels. Only basic DNA sequences were used in the experiments together with theoretical calculation to show the change in the energy level switching related to the Fermi level of the contact electrodes. I propose different quasiperiodic DNA sequences can be used to undergo similar tests and experiments. The Electromodulation spectra of Anti-symmetric (Thue–Morse) superlattice with different modulating voltage are shown in Figure 9 of the appendix. The different level of carrier states in the unconfined region are increased when the modulating voltage are increased. Along with similar research goal, the studies of Anti-symmetric sequences of DNA with different EC gate voltage can provide more insight into the CT behaviours. In particular, the quantum computer need to use some sophisticated nanostructures to carry out the quantum computing. Controlling the switching of different levels effectively with less energy consumption requires high CT rate in the unconfined region of quantum wells. Therefore, the proposed Anti-symmetric sequence could be a candidate that meets the demand of the quantum computing.

The Fibonacci sequences and related ratio were also found to have interesting connection with the nucleotide frequencies in single-stranded DNA of human genome.**[22–24]** Investigation of the Golden Ratio 1.618 and quasi-periodic sequences in nature can gain insight into some of the physical characteristic of quasi-periodic systems. Although the studies of Quantum walk on Protein-DNA target search**[25]** and aperiodic space-inhomogeneous system can provide localization properties and enhancement of entanglement**[26]** in quantum systems, the Quantum Well model presented in this study can facilitate and complement the investigation in DNA interaction and nanoelectronics.

## 6. Conclusion

In conclusion, we have performed experimental and theoretical studies of the unconfined state in the quasi-periodic sequences of Fibonacci and Anti-symmetric samples for DNA nanostructures and superlattices. When compared with the periodic and random samples, the Fibonacci and Anti-symmetric sample enhances the unconfined transitions by an order of magnitude in strength (Figure 8 and 9 in appendix). The experimental results were in agreement with the Quantum Well model simulation and provide foundation to expand the application to DNA nanostructure, which has dimension of about an order magnitude less than those of superlattice. The same Quantum Well model and computation method were used for the quasi-periodic sequences of Fibonacci and Anti-symmetric samples for DNA nanostructures. The effective mass value of 5 was selected for the carrier (electron or hole) transfer in the simulations as in Figure 4 – 6. When changing the effective mass values, the simulation results of transmission peaks can be shifted. More accurate determination of the effective mass value in DNA nanostructure will improve the simulation results when compared with the experimental results. The results published by the study of Thermoelectric effect and its dependence on molecular length and sequence in single DNA molecules**[1]** were used to compare with the simulations by the quantum well model. The results show agreement and can provide new insight into charge transfer and transport in DNA nanostructure with different types of sequences. Further experiments and analysis can be carried out to gain more insight and understanding concerning the characteristic of the Fibonacci and Anti-Symmetric sequences in DNA.

The cellular diagnostic mechanism**[20,27]** using the DNA CT assists the scanning of the genome to localise the damage and mutational sites that help to bring in the DNA repair protein. There will be many benefits, such as preventing uncontrolled mutations and improving cancer treatment, in studying the quantum mechanical effects and efficient communication across the DNA. Meanwhile, the experience and knowledge of the theoretical analysis, computer simulation and experimental results of quasi-periodic quantum systems can be applied for the development of the nanoelectronics circuits for quantum computing and other quantum devices.

## 7 Acknowledgments

The author like to thank M. Graf, D. Broido, P. Bakshi and K. Kempa for helpful discussion. I also like to thank T. Moustakas and T. Chiu for growing high-quality superlattices in the experiments. I appreciate Kayi Chan, Manni Mo and Breanna Tai for their comments and feedback.

## 9. Appendix

We review some of the previous work**[13]** for the experimental techniques of photoreflectance spectroscopy that were used to investigate the optical transitions between the unconfined states in different superlattice systems. The experimental results are compared with transition energies computed by the Quantum Well Transfer Matrix Method (TMM) to get the transmission coefficients in different energy states. When the transmission coefficients peaks are narrow and approaching to 1, the charge transfer of the carriers are also increased. The same computational method was used to calculate the functionality of the charge transfer in the DNA systems.

Three different superlattice samples[28] were grown on GaAs substrates in the (100) orientation using a VG-V80H MBE system under similar growth conditions:

i. Periodic, with alternating layers of AlGaAs (A) and GaAs(B).
ii. Quasiperiodic, with layers deposited according to the Fibonacci sequence.
iii. Random, with layers A and B selected by a random-number generator.

In all samples the Al mole fraction, x, =0.3; the AlGaAs (A) barrier width =40Å; and the GaAs (B) well width =28.3Å. In each instance, the total of barrier and well widths is made up of 600 layers (either A or B) resulting in a film thickness of approximately 2μm. The periodic sample was prepared to serve as a comparison with the other samples.

In order to build relations between the charge transfer of a superlattice semiconductor and DNA, a one-dimensional sequence with 1 or 0 will be used to simulate the sequence of a quantum well and barrier respectively. The superlattice samples were prepared on GaAs (undoped) substrates using a MBE system with the basic unit length of an AlGaAs barrier (0) = 40Å, and that of a GaAs well (1) = 28.3Å. Contrastingly, the helical chains of nucleotides in DNA are bounded to each other by hydrogen bonds that coil into tight loops and to form different shapes of polymers.

The existence of localized unconfined energy states in multiple periodic GaAs-AlGaAs quantum wells have been observed by photoluminescence spectra[29] and by photoreflectance spectroscopy[30]. Several material and system parameters were found to influence the transition probability within the unconfined regions. In the periodic GaAs-AlGaAs system, for example, wider barrier widths[29] and smaller Al mole fraction[30] have been found to enhance these transitions. Similar studies, however, of the unconfined energy states in the quasi-periodic and random quantum well structures have not received much attention[31]. The optical transitions involving the unconfined states of GaAs-AlGaAs in quasi-periodic, periodic and random superlattices are compared. In the random superlattice, the well widths and/or the barrier widths are varied according to an arbitrary random pattern. In contrast, the Fibonacci superlattice[32] is made up of an arrangement of layers of type A and type B following the Fibonacci sequence S1=A, S2=B, S3=AB, S4=BAB,…, Si=Si-2 Si-1. This leads to a quasiperiodic sequence of wells and barriers. Its band structure and wave-function localization in the confined states region have been shown to be dependent on the order of quasiperiodic modulation.

Photoreflectance is an optical technique for investigating the material and electronic properties of thin films. Photoreflectance measures the change in reflectivity of a sample in response to the application of an amplitude modulated light beam. In general, a photoreflectometer consists of an intensity modulated “pump” light beam used to modulate the reflectivity of the sample, a second “probe” light beam used to measure the reflectance of the sample, an optical system for directing the pump and probe beams to the sample, and for directing the reflected probe light onto a photodetector, and a signal processor to record the differential reflectance. The pump light is typically modulated at a known frequency so that a lock-in amplifier may be used to suppress unwanted noise, resulting in the ability to detect reflectance changes at the ppm level.

As shown in figure 8a, there are only weak transitions in the unconfined region for the periodic sample. In figure 8b, there are 2 conspicuous features appeared in the unconfined region for the Fibonacci superlattices. In figure 8c, there are overlapping features with the unconfined transition magnitude lie between those of periodic and Fibonacci superlattices for the random superlattices.

The corresponding computed results[13] were compared with the experimental results They agree well with the experimental results such that

i. Periodic sample: There is a wide gap appeared between the top of the barrier and the first unconfined energy band above the barrier.
ii. Fibonacci sample: There are distinct features of the unconfined states appeared very close to the barrier.
iii. Random samples: There is overlapping features of unconfined states appeared above the barrier.

The strong signals obtained in the unconfined region of the Fibonacci sample observed in the photoreflectance spectra matched with the theoretical calculation. The narrow energy bandwidth corresponding to the unconfined transition implies a longer time spending in the well because of the Uncertainty Principle. The longer time the electron spends in the well, the higher probabilities for the electron to be captured by the well. Therefore, the transition strength is enhanced. The Anderson localization[33] effect shows that the wavefunctions form the wave packet to become a localized state. The carrier transport efficiency decreases along the growth axis of purposely disordered or random GaAs/GaAlAs superlattices. On the other hand, the carrier transport efficiency increases when the bandwidth of the transmission peaks is distinct and narrow in the simulation of the Fibonacci sample. The large signals of “A” and “B” in figure 8b validate the theory and simulation of this study. When compared with the periodic and random samples, the Fibonacci sample enhances the unconfined transitions by an order of magnitude in strength.

**Figure 8.**
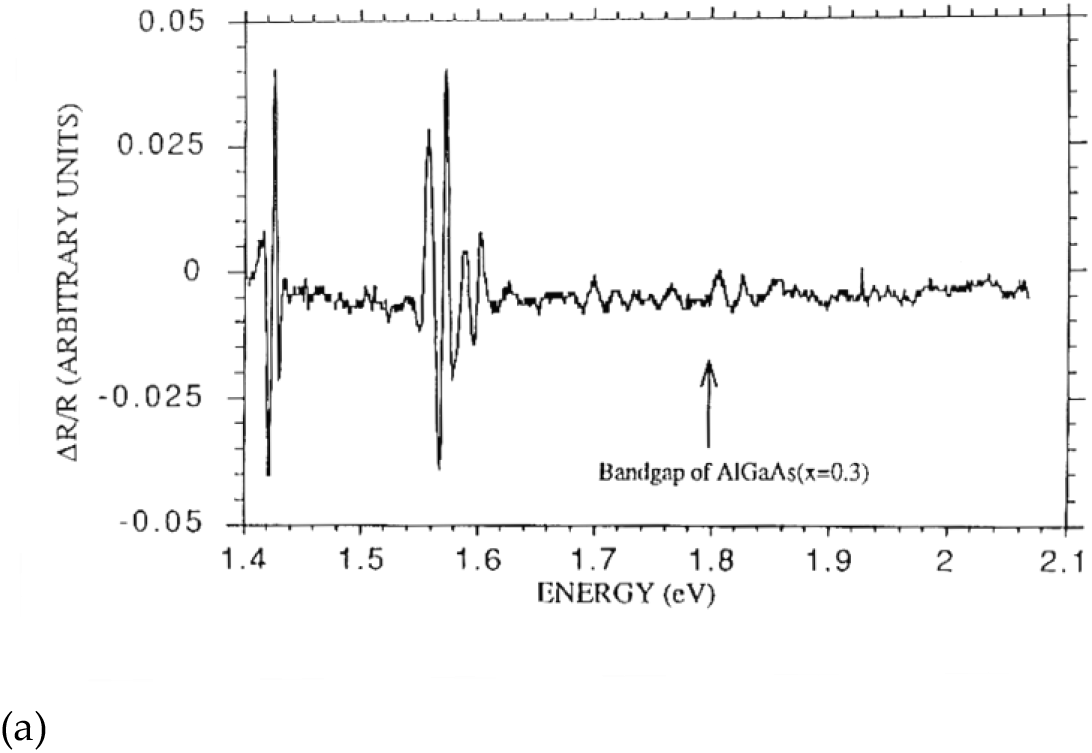
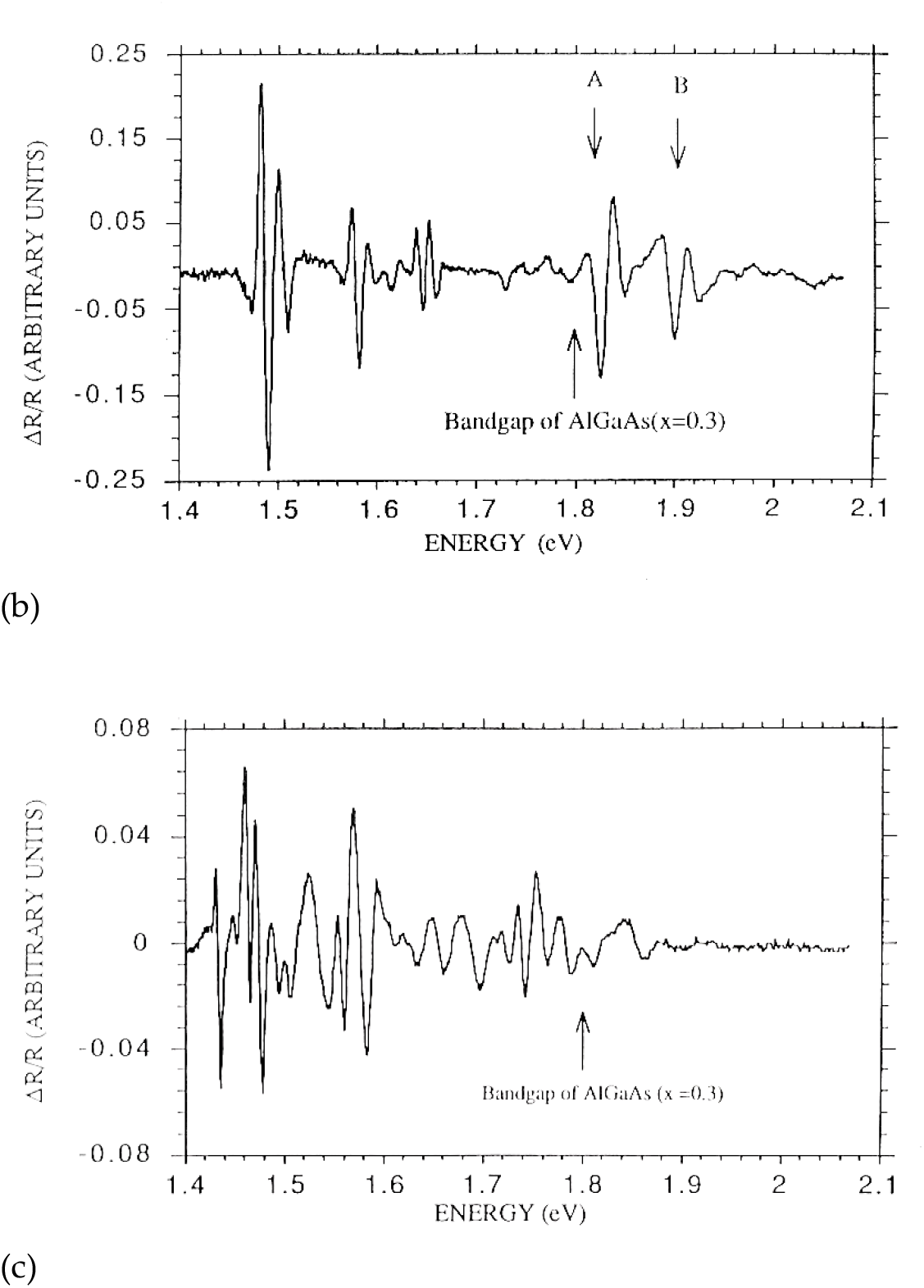
Photoreflectance spectra of (a) periodic (b) Fibonacci (c) random superlattice at room temperature.

**Figure 9.**
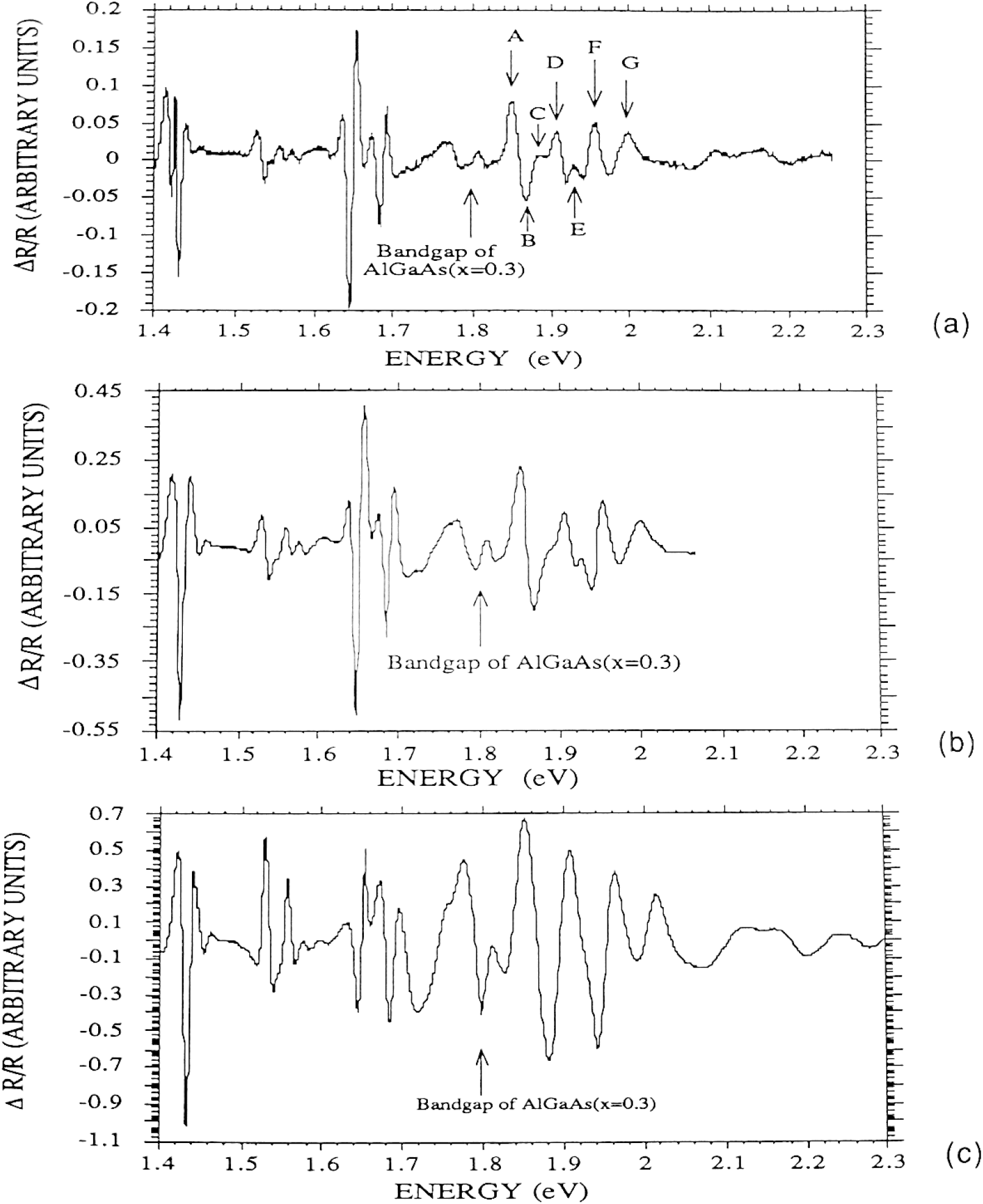
Electromodulation spectra of Anti-symmetric (Thue–Morse) superlattice with modulating voltage at (a) 4V_p-p_ (b) 10V_p-p_ (c) 20V_p-p_. The measurements were performed at room temperature and the bias was at 0V bias.

The transmission peak is sensitive to the number of layers considered in the calculation. For quasi-periodic superlattice (Fibonacci sequence, table 2), the general patterns remain the same when the number of boundaries are changed. In particular, the superlattice with 44 boundaries has unity transmission peaks while the others are not quite to have these unity transmission peaks. The resolution of the calculated transmission coefficients is the same and equal to 0.1meV. We find that the 44 boundaries of the Fibonacci superlattice correspond to the first symmetric sequence, i.e. the half left sequence is the image of the half right sequence. This is the criteria for generating a unity transmission peak. At the other boundaries, the half left sequences do not form a perfect mirror image of the corresponding half right sequence. There exist one or more units in the sequence that does not match the image requirement. These result in generating the transmission peaks less than unity besides the slight shift of their energy position. These can be further supported by the transmission plot in the “anti-symmetric” superlattice (table 3). The sequence that forms the superlattice actually meets the image matching requirement at layers 1 to 64. This result in a unity transmission peaks. Another image matching occur at layers 1 to 256 (172 boundaries) which give rise to unity transmission peaks of the simulations (Figure 6b). These are one of the characteristic of quasi-periodic superlattice that cannot be found in the random superlattice. This preservation of transmission coefficient of quasi-periodic superlattice is especially an advantage for the study of optical transition. The corresponding position of the transitions remain the same even though the penetration depth of the probe and pump light are changed, i.e. the number of layers are changed. Therefore, this give rise to consistent transition energy, in which the combined effect can increase the transition strength.

The unconfined state transition of the superlattices belong to the band to band transition. Hence, the application of large electric field can generate the Franz-Keldysh effect[34]. The depletion layer of the superlattice is estimated to be in the order of 10,000Å. The strength of the electric field is in the order of 10^6^ V/m when 10V is applied across the sample. This large electric field can result in Franz-Keldysh effect. As shown in fig. 9 (a-c), the relative amplitude of the unconfined transition signal to GaAs signal of the Anti-symmetric superlattice increase almost two-fold when the modulation voltage increases from 4Vp-p to 20Vp-p. There are six peaks in the unconfined region at 4Vp-p. While the voltage increases, two of the peaks shrink and the total peaks end up in four large peaks at 20Vp-p. The unconfined state transition strengths were enhanced for the quasi-periodic sample (Anti-symmetric superlattice) when compared with samples having periodic and random sequences. The experimental results of the measured extended transition energies agree qualitatively with theoretical Transfer Matrix (TMM) calculations. Further experiments and analysis will be performed to gain more insight and understanding concerning the characteristics of the Anti-symmetric sample with respect to superlattice and DNA nanostructure.

